# Transposable elements contribute to the spatiotemporal microRNA landscape in human brain development

**DOI:** 10.1101/2022.01.14.476326

**Authors:** Christopher J. Playfoot, Shaoline Sheppard, Evarist Planet, Didier Trono

**Affiliations:** School of Life Sciences, Ecole Polytechnique Fédérale de Lausanne (EPFL), 1015 Lausanne, Switzerland

## Abstract

Transposable elements (TEs) contribute to the evolution of gene regulatory networks and are dynamically expressed throughout human brain development and disease. One gene regulatory mechanism influenced by TEs is the miRNA system of post-transcriptional control. miRNA sequences frequently overlap TE loci and this miRNA expression landscape is crucial for control of gene expression in adult brain and different cellular contexts. Despite this, a thorough investigation of the spatiotemporal expression of TE-embedded miRNAs in human brain development is lacking. Here, we identify a spatiotemporally dynamic TE-embedded miRNA expression landscape between childhood and adolescent stages of human brain development. These miRNAs sometimes arise from two apposed TEs of the same subfamily, such as for L2 or MIR elements, but in the majority of cases stem from solo TEs. They give rise to *in silico* predicted high-confidence pre-miRNA hairpin structures, likely represent functional miRNAs and have predicted genic targets associated with neurogenesis. TE-embedded miRNA expression is distinct in the cerebellum when compared to other brain regions, as has previously been described for gene and TE expression. Furthermore, we detect expression of previously non-annotated TE-embedded miRNAs throughout human brain development, suggestive of a previously undetected miRNA control network. Together, as with non-TE-embedded miRNAs, TE-embedded sequences give rise to spatiotemporally dynamic miRNA expression networks, the implications of which for human brain development constitute extensive avenues of future experimental research. To facilitate interactive exploration of these spatiotemporal miRNA expression dynamics, we provide the “Brain miRTExplorer” web application freely accessible for the community.

## Introduction

Transposable elements (TEs) account for around half of the human genome and have contributed to the evolution of gene regulatory networks. The majority of TEs have lost their capacity to ‘copy and paste’ to new locations around the genome, instead being co-opted by the host organism to perform a plethora of regulatory homeostatic functions during normal development (Elbarbary et al. 2016; Chuong et al. 2017). One post-transcriptional regulatory mechanism in which TE-embedded sequences have been co-opted is the microRNA (miRNA) system (Smalheiser and Torvik 2005; Piriyapongsa et al. 2007; Roberts et al. 2014). Computational and experimental studies have shown different classes of TEs (LINE, SINE and LTR) can act as functional sources of miRNA in different cellular models. However, limited information exists for primary tissues, especially for tightly regulated spatiotemporal developmental processes such as human brain development (Piriyapongsa et al. 2007; Piriyapongsa and Jordan 2007; Ding et al. 2010; Frankel et al. 2014; Roberts et al. 2014; Spengler et al. 2014; Petri et al. 2019). Recent studies in a small number of adult brains have highlighted the roles of TE-embedded miRNAs from the L2 family. These are functional in neurotypical adult brains and are differentially expressed in glioblastoma (Skalsky and Cullen 2011; Petri et al. 2019). Furthermore, miRNAs have critical roles in mammalian neuronal homeostasis, highlighting the fundamental nature of miRNAs in neurogenesis, alongside diverse roles in neurological disease and human evolution (Cao et al. 2007; Somel et al. 2011; Qureshi and Mehler 2012; Petri et al. 2014; Topol et al. 2016; Sambandan et al. 2017; Juźwik et al. 2019; Woods and Van Vactor 2021). miRNAs are spatially and temporally expressed in the developing human brain from birth to adolescence, however the contribution of TE-embedded sequences to this process has never been investigated (Ziats and Rennert 2014). Indeed, the years proceeding birth and throughout childhood represent a crucial window in human brain development, characterized by extensive changes in size, cellular composition and functional processes such as synaptogenesis, myelination and synaptic pruning (Silbereis et al. 2016; Dyck and Morrow 2017). We therefore aimed to determine the prevalence of spatiotemporally expressed, annotated TE-embedded miRNAs in the developing human brain by re-analysis of small RNA-seq data available from the BrainSpan Atlas of the Developing Human Brain from one year old to 19-year-old brains (Miller et al. 2014; Li et al. 2018). We computationally uncover dynamic spatiotemporal expression of numerous annotated TE-embedded miRNAs and a small number of previously undetected novel putative TE-embedded miRNAs, suggesting TE-sequence co-option as miRNAs may play a role in this important neurodevelopmental window. We provide the “Brain miRTExplorer” web application to facilitate interactive exploration of both annotated TE-embedded and non-TE-embedded miRNA spatiotemporal expression data, freely accessible for the community at https://tronoapps.epfl.ch/BrainmiRTExplorer/.

## Results

### TEs contribute to the annotated miRNA transcriptional landscape in the human brain

To determine spatiotemporal, small RNA expression in postnatal human brain development, we analyzed small RNA-seq data from 174 samples from one year to 20 years of age, encompassing 16 different brain regions, from 16 donors (9 male and 7 female) available through the BrainSpan Atlas of the Developing Human Brain (Supplemental Fig. S1) (Miller et al. 2014; Li et al. 2018). To enrich for different small RNA moieties, we separated sequencing reads into lengths of 18 - 25bp, 26 – 37bp and 38 – 50bp and intersected with Ensembl annotations, miRBase, the GtRNAdb database and our modified merged TE RepeatMasker data set (Kozomara and Griffiths-Jones 2014; Chan and Lowe 2016; Pontis et al. 2019; Turelli et al. 2020; Yates et al. 2019; Playfoot et al. 2021). As expected, the different read lengths enriched for annotated miRNAs, tRNAs and snoRNAs respectively (Fig. 1A; Supplemental Fig. S2). By retaining the miRNA derived 18 – 25bp reads, we detected the expression of 543/1871 annotated miRNAs (Fig. 1B; Supplemental Table S1). To determine the overlap of annotated miRNAs with TEs, we intersected their genomic coordinates with those from our curated RepeatMasker data set (Turelli et al. 2020; Playfoot et al. 2021). 17% of annotated miRNAs were derived from TEs, in either sense and antisense orientation to the miRNA and belonged to all known classes of elements with representatives from various subfamilies and evolutionary ages (Fig. 1B, C; Supplemental Table S1). L2 family members of 105 - 177 million-year-old (MYO) contributed the most to annotated expressed miRNAs in the child and adolescent brain (Fig. 1C), with detection of all L2-embedded, annotated miRNAs previously noted in adult brain and glioblastoma (Piriyapongsa et al. 2007; Petri et al. 2019), pointing to their likely roles in earlier stages of brain development (Supplemental Table S1). The previously described 43.2 MYO MADE1 elements and the 177 MYO MIR family elements also heavily contributed to expressed miRNAs (Fig. 1C) (Piriyapongsa and Jordan 2007; Shao et al. 2010; Borchert et al. 2011; Spengler et al. 2014).

**Figure 1.**
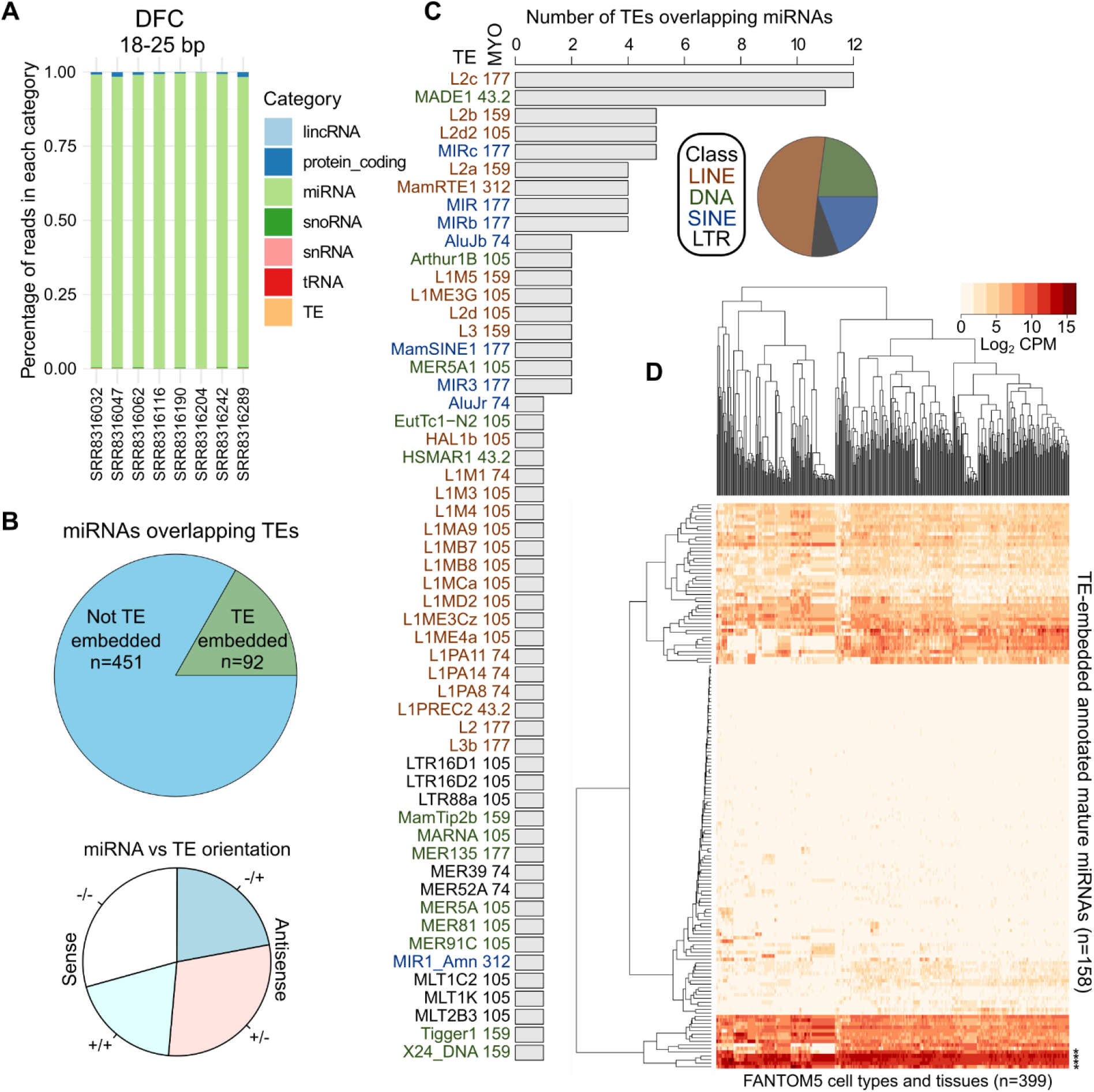
TEs contribute to annotated miRNAs in the child and adolescent human brain. (*A*) Stacked bar chart indicating the percentage of 18-25bp reads overlapping different annotated genomic features for samples from the dorsolateral frontal cortex (DFC). If a TE overlaps an annotated feature (miRNA, tRNA etc.) the feature takes preference. (*B*) Pie charts indicating the number of miRBase annotated miRNAs overlapping at least one TE (*top*), and their relative orientations (*bottom*). (*C*) Bar chart indicating the number of TEs overlapping miRBase annotated miRNAs and their class and age in million years old (MYO). (*D*) Expression in log2 counts per million (CPM) of mature TE-embedded miRNA in 399 cell types and tissues from FANTOM5 (De Rie et al. 2017). * denotes miRNAs highlighted in the text.

We next aimed to determine if TE-embedded miRNAs were produced in other cell types and tissues by analyzing miRNA expression data from 399 human samples (De Rie et al. 2017). Mature TE-embedded miRNAs were broadly expressed in the majority of cell types, with relatively ubiquitous, high levels for MIRc-embedded hsa-miR-378a-3p, L2d2-embedded hsa-miR-28-3p, L2c-embedded hsa-miR-151a-3p/5p and MamRTE1-embedded hsa-miR-130a-3p and lower expression for other TE-embedded miRNAs (Fig. 1D). Together, these data indicate that a multitude of TE-embedded miRNAs are broadly expressed in the child and adolescent human brain, with appreciable expression in other cell types.

### TE-embedded miRNAs exhibit spatiotemporal expression patterns

To investigate the temporal dynamics of TE-embedded miRNAs in brain development, we compared their expression from childhood (1 to 5 years) to adolescence (9 to 20 years) (Supplemental Fig. S1). We initially combined samples of forebrain (FB) origin, representing 124 samples from 16 donors, with 66 and 58 samples representing childhood and adolescence respectively (Supplemental Fig. S1). 16% and 5.5% of TE-embedded miRNAs were significantly more highly expressed in childhood or adolescence respectively, whilst 78% were continually expressed (Fig. 2A). Differentially expressed miRNAs, again represented a suite of TE subfamilies and evolutionary ages (Supplemental Table S1). There was no difference in the ages of differentially expressed or continual TE-embedded miRNAs (Supplemental Fig. S3A). Non-TE-embedded miRNAs exhibited similar expression patterns, indicating that temporal expression is not restricted to TE-embedded miRNAs but is a broad feature of this class of post-transcriptional regulators (Ziats and Rennert 2014) (Supplemental Fig. S3B).

**Figure 2.**
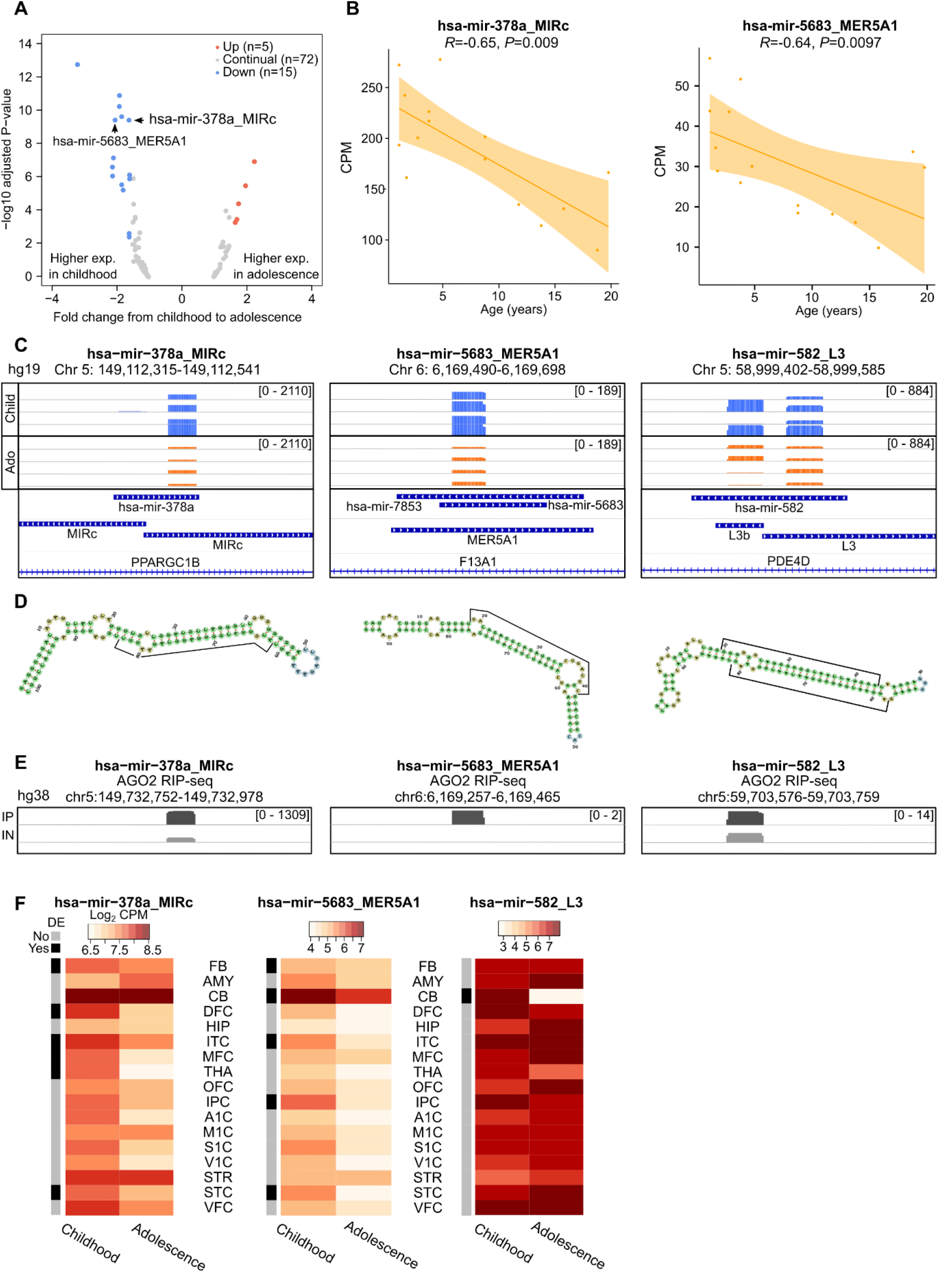
TE-embedded miRNAs are temporally expressed between child and adolescent human brains. (*A*) Volcano plot highlighting TE-embedded miRNAs significantly differentially expressed in FB (adjusted *P-value ≤ 0.05, 1.5-fold change*). (*B*) Dot plots showing the correlation of expression and age for specific TE-embedded miRNAs. Shaded area represents the variance. (*C*) Integrated genome viewer (IGV) visualization of four childhood (blue) DFC BAM files and four adolescent (orange) DFC BAM files, alongside miRBase annotation, TE annotation and gene annotations for hg19. Read count is shown within square brackets. (*D*) miRNA hairpin schematics from miRNAfold (Tempel and Tahi 2012; Tav et al. 2016) for the DNA sequences in *C*. Each hairpin structure exhibits 90% of verified miRNA hairpin features as previously defined (Tempel and Tahi 2012; Tav et al. 2016). 22bp peak sequences are highlighted by the black bars on arms of the hairpin. (*E*) IGV visualization of the corresponding region of *C* but in hg38 for AGO2-RIPseq in human embryonic stem cell derived neurons for AGO2 immunoprecipitated (IP) and input (IN) samples (Petri et al. 2019). (*F*) Heatmaps showing regional expression in log2 counts per million (CPM), alongside differential expression results (black and grey bar). Region abbreviations are defined in Supplemental Fig. S1.

In order to confirm our differential expression results, we next matched the expression of TE-embedded miRNAs in the FB with donor age. Of the 20 differentially expressed TE-embedded miRNAs, 12 also exhibited significant correlations or anti-correlations with this parameter (Supplemental Fig. S3C; Supplemental Table S2).

One of the most significantly differentially expressed TE-embedded miRNAs in the FB was the cancer- and cell proliferation-associated hsa-mir-378a, which displayed higher expression in childhood and a significant anti-correlation with donor age (Li et al. 2015; Velazquez-Torres et al. 2018; Guo et al. 2019)(Fig. 2A, B *left* & C *left*). This miRNA is embedded in two intronic, MIRc elements arranged in opposite orientations, facilitating high confidence pre-miRNA hairpin precursor formation as determined by *in silico* miRNA folding analyses to detect hairpins with 90% of verified miRNA hairpin features (Tempel and Tahi 2012; Tav et al. 2016)(Fig. 2D *left*). The glycolysis-, cancer- and cell proliferation-associated hsa-mir-5683 was also significantly more expressed in childhood, with a significant anti-correlation with donor age, however was embedded in a solo 105 MYO MER5A1 element which also facilitated pre-miRNA hairpin formation (Miao et al. 2020; Rong et al. 2020) (Fig. 2A, B *right*, C *middle* & D *middle*). One TE-embedded miRNA which was continually expressed in childhood and adolescent brains was the cancer- and neuron-associated hsa-mir-582, embedded in two apposed L3 elements, again leading to an *in silico* predicted high-confidence precursor hairpin structure (Fang et al. 2015; Zhang et al. 2015; Ding et al. 2019) (Fig. 2C *right* & D *right*). Indeed, 18/92 miRNAs overlapped at least two TEs, with varying genomic orientations (Supplemental Table S3), although the majority of expressed TE-embedded miRNAs overlapped only one TE.

We next aimed to confirm our detection of these three TE-embedded miRNAs by using an independent Argonaute2 RNA-immunoprecipitation sequencing (AGO2 RIP-seq) data set from human embryonic stem cell-derived neurons, independently mapped to hg38 (Petri et al. 2019). AGO2 directly binds to mature processed miRNAs for incorporation into the RISC complex for targeting of mRNA (Kobayashi and Tomari 2016; Michlewski and Cáceres 2019). AGO2-bound elements are thus likely to represent *bona fide* miRNAs rather than mere degradation products. Enrichment of reads in the AGO2-immunoprecipitation sample was observed compared to the input sample, with peaks residing over exactly the same sequences as in our hg19-mapped data (Fig. 2E).

### Different regions exhibit diverse miRNA temporal expression patterns

The temporal TE and gene expression profile of the human brain varies by region, notably with the cerebellum (CB) displaying a different transposcriptional and transcriptional landscape when compared to FB (Playfoot et al. 2021). We therefore next determined the temporal expression profile of miRNAs in childhood and adolescence in different individual brain regions (Fig. 2F, Supplemental Table S1). hsa-mir-378a MIRc exhibited significantly higher expression in childhood not only in combined FB samples, but also in individual FB regions such as the dorsolateral prefrontal cortex (DFC), the inferior temporal cortex (ITC), the medial prefrontal cortex (MFC) and the superior temporal cortex (STC), along with non-FB regions such as the mediodorsal nucleus of the thalamus (THA) (Fig. 2F *left*). Similarly, hsa-mir-5683 MER5A1 had significantly higher expression in childhood versus adolescence in the FB combined, along with other individual FB regions, but also in the CB (Fig. 2F *middle*). In both instances, expression in the CB was higher than for any other individual region. In contrast, the L3-embedded hsa-mir-582 exhibited continually high expression across childhood and adolescence for all regions, except the CB where hsa-mir-582 L3 expression was largely absent in adolescence and restricted to childhood (Fig. 2F). This provides a striking example of spatiotemporal control of the miRNA transcriptional landscape. Overall, these data demonstrate that the TE-embedded miRNA transcriptional landscape exhibits diverse spatiotemporal dynamics, with sometimes overt differences between childhood and adolescence for FB and non-FB regions.

### TE-embedded miRNAs are spatially expressed

Due to the temporal nature of miRNA expression in multiple brain regions, we next aimed to determine spatial differences in TE-embedded miRNA expression, regardless of age. We performed 120 differential expression analyses, comparing each region to each other independent region. Of these comparisons, the region with the largest number of differentially expressed TE-embedded miRNAs was consistently the CB (Fig. 3A). Of the top 30 comparisons with the highest number of differentially expressed TE-embedded miRNAs, the CB was responsible for half (Fig. 3A). The CB vs the hippocampus (HIP) had the highest number of differentially expressed TE-embedded miRNAs, followed by the CB vs striatum (STR), amygdala (AMY) and many regions of the FB (Fig. 3A). These data suggest that the CB exhibits not only different TE and gene expression compared to other brain regions as previously described (Playfoot et al. 2021), but also differences in TE-embedded miRNA expression.

**Figure 3.**
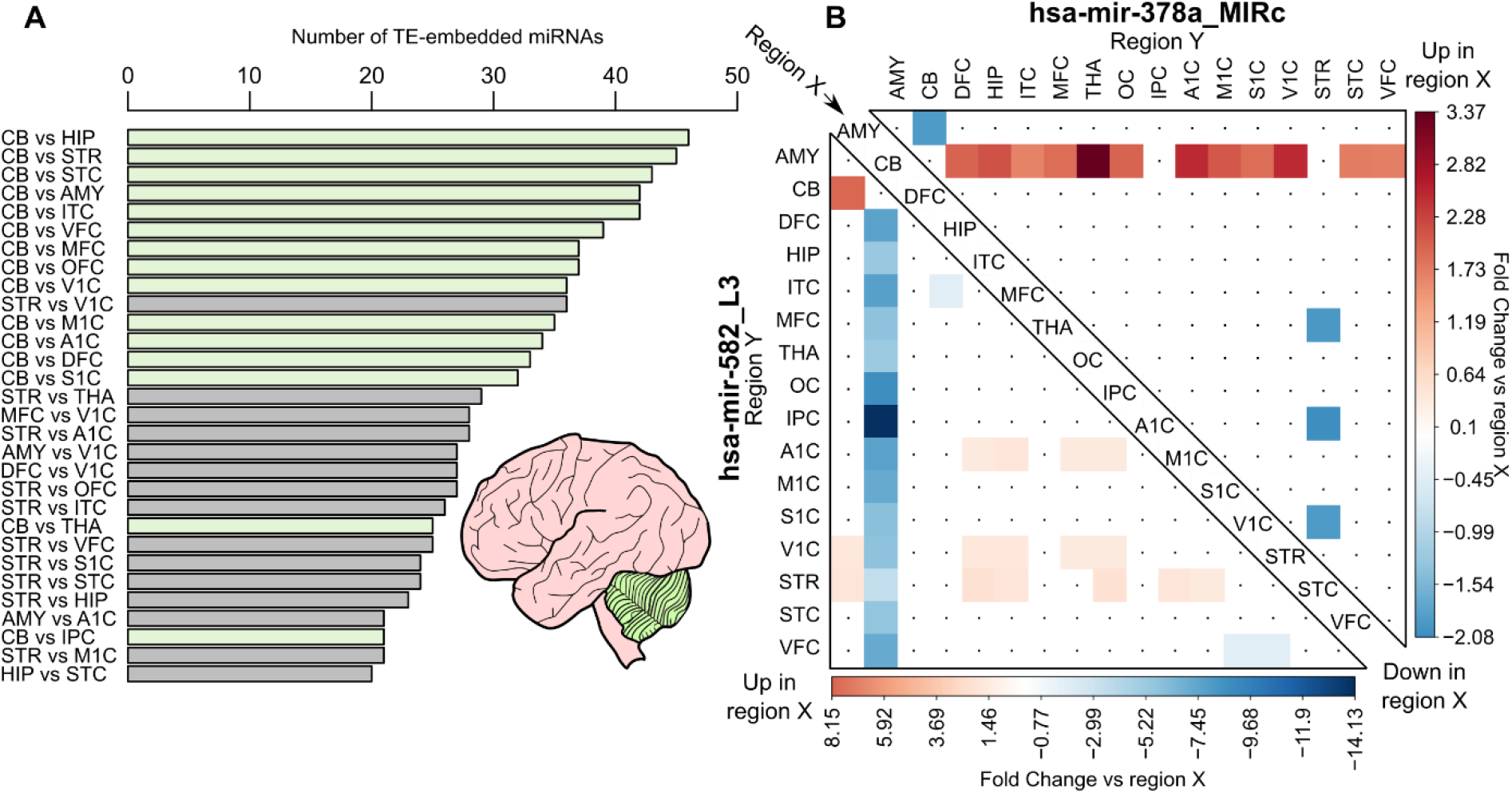
TE-embedded miRNAs exhibit spatial expression with major differences in the cerebellum. (*A*) Bar chart showing the number of differentially expressed TE-embedded miRNAs per regional comparisons (*P-value ≤ 0.05*, 1.5-fold change up or down). Only the top 30 comparisons are shown. (*B*) Heatmap comparing the fold change of region X (*center diagonal*) to region Y (*left and top*) for two TE-embedded miRNA loci described in Fig. 2. Only regions with significant fold changes are colored (*P-value ≤* 0.05, 1.5-fold change).

As hsa-mir-378a MIRc exhibited distinct temporal expression (Fig. 2) we next assessed its potential spatial expression. Indeed, the CB exhibited significantly higher expression of hsa-mir-378a MIRc when compared to most other regions (Fig. 3B). Conversely, hsa-mir-582 L3 exhibited significantly lower expression in the CB compared to all other regions, suggestive of diverse regulatory control of different miRNAs (Fig. 3B). A multitude of other examples of spatial miRNA expression suggests widespread spatial regulation of not only TE-embedded miRNAs, but also non-TE-embedded miRNAs. These dynamics can be interactively explored for all miRNAs with our Brain miRTExplorer application.

### TE-embedded miRNAs target neurogenesis-associated genes

In order to determine possible functional relevance, we extracted predicted genic targets of TE-embedded miRNAs from the TargetScan database (Supplemental Table S4) (Agarwal et al. 2015; McGeary et al. 2019). We specifically focused on conserved TE-embedded miRNAs and conserved genic targets (Agarwal et al. 2015; McGeary et al. 2019). Using this stringent list, gene ontology (GO) biological process analysis indicated that many target genes of TE-embedded miRNAs are enriched in neurogenesis-associated functions. For example, the L2c-embedded hsa-mir-374b targets all four genes involved in striatal medium spiny neuron differentiation (GO:0021773) and three out of four genes associated with glial cell fate specification (GO:0021780) and oligodendrocyte cell fate specification (GO:0021778). Similarly, the L2b-embedded hsa-mir-493 was enriched in positive regulation of synaptic vesicle exocytosis (GO:2000302) and neurotransmitter receptor transport to plasma membrane (GO:0098877), among others. These analyses also revealed significant enrichments in GO cell component terms for hsa-mir-493 such as NMDA selective glutamate receptor complex (GO:0017146), glial cell projection (GO:0097386) and integral component of postsynaptic specialization membrane (GO:0099060), among many other neurogenesis associated terms. For the aforementioned L3-embedded hsa-mir-582, enrichment in terms such as astrocyte end-foot (GO:0097450) and main axon (GO:0044304) was detected. The diverse GO enrichments detected for different TE-embedded miRNA predicted targets highlight the specialized miRNA target networks in human neurogenesis.

### TEs contribute to novel putative miRNAs

Most studies rely on mapping small RNA-seq reads directly to miRNA annotations provided in miRBase. As a large proportion of annotated miRNAs are embedded in TEs, we reasoned that other TE loci could be contributing to previously undetected, novel miRNAs expressed in the brain. We therefore further investigated our unbiased, unique mapping to the whole genome used for detection of annotated TE-embedded miRNAs. To ensure robustness and to limit false positives we used our custom RepeatMasker annotation (Turelli et al. 2020; Playfoot et al. 2021), alongside manual curation by inspecting BAM files from childhood and adolescent samples of the DFC to detect a characteristic 22bp peak. We next focused on two of the most robust candidates. The first was embedded in two apposed head-to-head, intronic MER3 elements and was confirmed with peaks detectable in the AGO2-RIPseq data, suggestive of processed miRNA (Fig. 4A & B *left*). Indeed, the 200bp sequence covering the miRNA locus facilitated *in silico* hairpin structure formation with 90% of verified features and the 22bp 3p and 5p peak sequences contributing to each arm of the hairpin (Fig. 4C *left*). The same was observed for a novel putative miRNA embedded in a single MER5A element (Fig. 4A, B & C *right*). To determine the evolutionary history of these two loci, we assessed the 22bp sequence using MULTIZ alignments. Indeed, the MER3-embedded miRNA is present in rhesus macaque but absent from mouse, whereas the MER5A element is present in rhesus macaque but with multiple mutations in the seed region in mouse (Fig. 4D). To determine their novelty, the 22bp sequence of these candidates were searched in miRBase and did not match any sequences. These two TE loci therefore represent robust, novel TE-embedded miRNAs, the function of which remains to be elucidated. Together, these data highlight the dynamic spatiotemporal nature of annotated and novel TE-embedded miRNAs in the developing human brain and provides scope to investigate the disease and functional relevance of TE sequence co-option as miRNAs throughout evolution.

**Figure 4.**
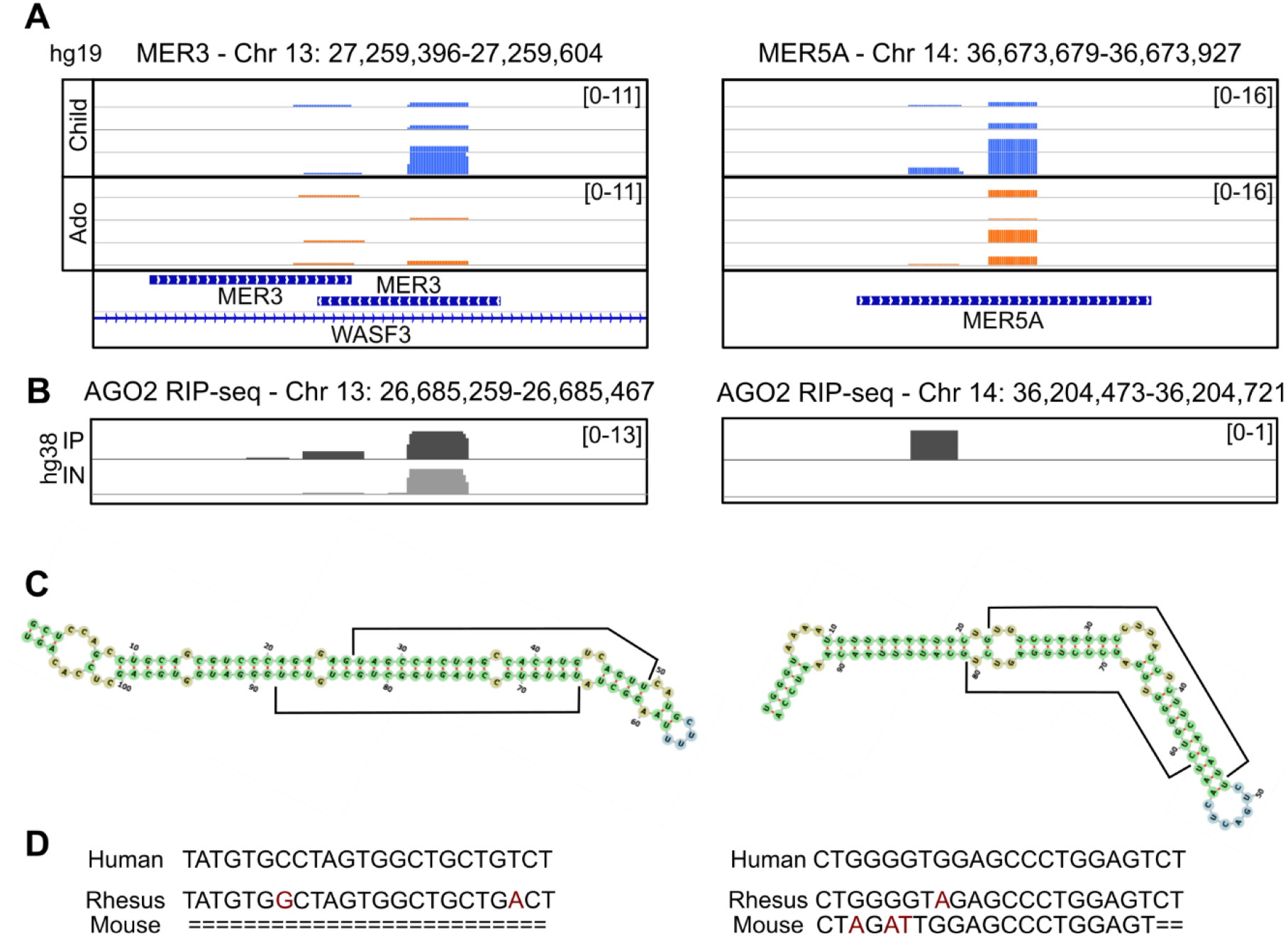
Novel, non-annotated TE-embedded miRNAs are present in child and adolescent brains. (*A*) IGV visualization of non-annotated TE-embedded miRNAs with classical 22bp peaks in four childhood (blue) DFC BAM files and four adolescent (orange) DFC BAM files, alongside TE and gene annotations for hg19. (*B*) IGV visualization of the corresponding region in *A* but in hg38 for AGO2-RIPseq in human embryonic stem cell derived neurons for AGO2 immunoprecipitated samples (IP) and input (IN) (Petri et al. 2019). (*C*) miRNA hairpin schematics from miRNAfold (Tav et al. 2016) for the DNA sequences in *A*. 22bp peaks are highlighted by the black bars on both arms of the hairpin. (*D*) MULTIZ alignment from the UCSC genome browser of the 22bp miRNA sequence beneath the largest peak in *A* (Blanchette et al. 2004; Navarro Gonzalez et al. 2021).

## Discussion

Human brain development is a dynamic and highly regulated spatiotemporal process, however the contribution of TEs to one mechanism of regulatory control, miRNAs, has never been formally investigated in this context. We show that the postnatal TE-embedded miRNA landscape is indeed spatially and temporally dynamic, with alterations in TE-embedded miRNA expression from childhood to adolescence, similar to non-TE-embedded miRNAs. Our previous work highlighted a distinct TE expression switch during late prenatal and early postnatal developmental timepoints, accompanied by coordinated reduction in expression of their controlling transcription factors, the KRAB-zinc finger proteins (KZFPs) (Playfoot et al. 2021). Furthermore, we determined spatiotemporal TE-mediated alternative promoter usage leading to novel mRNA transcript isoforms, indicative of direct TE-dependent transcriptional innovation (Playfoot et al., 2021). Here, we expand the role of TEs in human brain development to that of miRNAs; a more indirect, but no less important method of transcriptional innovation.

One critical limitation of our study is the restriction to postnatal timepoints. As major gene and TE expression changes occur during prenatal to postnatal transitional stages, future work should aim to generate small RNA-seq data covering the whole timeframe of human brain development. miRNAs were previously demonstrated to play critical roles in mouse prenatal brain development (Petri et al. 2014), and we found here that many human TE-embedded miRNAs were more highly expressed in childhood when compared to adolescence. As many neurological disorders appear to have origins in early development (Short and Baram 2019), it would be imperative to investigate both TE-embedded and non-TE-embedded miRNA expression at prenatal stages. To date, the limited number of human studies aiming to address this point were restricted by sample number, developmental stages and regions (Nowakowski et al. 2018). Whilst advances in human embryonic stem cell differentiation protocols have enabled *in vitro* study of different neurological cell types, these lack the wide diversity present in tissue samples.

The detection of novel, non-annotated TE-embedded miRNAs is suggestive of a previously undetected TE-originating miRNA landscape. The volume of data assessed may have allowed the detection of these, however computational limitations of using only uniquely mapping reads is especially acute for young, more homogenous TE subfamilies which have accumulated less mutations. Future work should experimentally assess putative, young repetitive TE-embedded miRNAs, as they have the potential to significantly expand the RNA-based regulome. Their repetitive nature likely facilitates post-transcriptional control of mRNA targets containing the same TE subfamilies in their 3’ UTRs, as has been shown for the annotated L2-embedded miRNAs (Petri et al. 2019). These results also suggest a multifactorial role for TEs, whereby some TEs give rise to mature miRNAs but are also targets of the miRNA microprocessor machinery themselves, thus acting to restrict their movement when the TEs remain retrotransposition competent (Heras et al. 2013, 2014).

In summary, the spatiotemporal expression of TE-embedded miRNAs from childhood to adolescence suggests a role for TEs in the fine tuning of transcriptional networks at the post-transcriptional level throughout human brain development. Although these dynamics are not restricted to TE-embedded miRNAs, these analyses provide a novel insight into a crucial understudied developmental window, as the role of TE-embedded miRNAs has only been previously investigated in adult or disease contexts.

## Materials and Methods

### Dataset download and preprocessing

Raw small RNA-seq FASTQ files from the BrainSpan Atlas of the Developing Human Brain (phs000755.v2.p1 provided by Dr. Nenad Sestan), were downloaded from the dbGaP-authorized access platform (Miller et al. 2014; Li et al. 2018) (Supplemental Acknowledgements). The reads were first trimmed to remove Illumina small RNA 3’ sequencing adapters (TGGAATTCTCGGGTGCCAAGG) using FLEXBAR (version 3.5.0) with parameters --adapter-trim-end RIGHT --min-read-length 18 (Dodt et al. 2012). Trimmed reads were then divided by read length ranges of 18-25 nucleotides, 26-37 nucleotides and 38-50 nucleotides. Reads were then mapped to the human hg19 genome (GRCh37.p5) using Bowtie (version 2.3.4.1) with parameter --very-sensitive-local (Langmead et al. 2009). Read counts on different genomic features were quantified using featureCounts (version 1.6.2 of the subread package) (Liao et al. 2014). Uniquely mapped reads were quantified with parameters -t exon -g gene_id -Q 10 and multimapped reads with parameters -M –fraction -t exon -g gene_id -Q 0. We used the parameters -s 1 and -s 2, to quantify sense and antisense reads respectively which were subsequently merged, keeping only the strand with the most reads. To confirm specific read lengths were enriching for specific RNA moieties, the annotation of snoRNA, snRNA, miscRNA, scRNA and genes from Ensembl (GRCh37.p5, release 100) were used. For miRNAs and tRNAs, miRBase version 20 (Kozomara and Griffiths-Jones 2014) and tRNA annotations from GtRNAdb (release 19) were used respectively (Chan and Lowe 2016). For repetitive sequences, a previously described in-house curated version of the RepeatMasker database was used (where fragmented LTR and internal segments belonging to a single integrant were merged) (Turelli et al. 2020; Playfoot et al. 2021). Exons of genes and TEs overlapping small RNAs in the same orientation, were removed using BEDTools intersect (version 2.27.1) with default parameters to prioritize reads falling on small RNAs (Quinlan and Hall 2010). To determine which expressed annotated miRNAs overlapped TEs, we used BEDTools to intersect the miRBase and our custom RepeatMasker merged TE annotations with a minimum of one base pair overlap. TE subfamily age estimates were obtained from DFAM (Hubley et al. 2016). BAM files were visualized using the Integrative Genome Viewer (Robinson et al. 2011).

### Filtering and normalization

Samples were sequenced with a read length of 51bp and samples with less than 1 million reads mapped were removed. Features where the sum of the counts over all the samples was lower than the total number of samples were removed. TEs overlapping gene exons were also removed using BEDTools closest (Quinlan and Hall 2010). Normalization for the sequencing depth was performed for all features on the sense and antisense with the TMM method as implemented in the R package limma (version 3.46.0) (Ritchie et al. 2015). The total number of mapped reads was used as library size.

### Differential expression analysis

Samples from one year to five years were considered as childhood and nine years to 20 years as adolescence (Supplemental Fig. S1). To perform the aggregated temporal FB differential expression, the following brain regions were considered as FB: Dorsolateral prefrontal cortex, Inferior temporal cortex, Medial prefrontal cortex, Orbital prefrontal cortex, Posterior inferior parietal cortex, Primary auditory (A1) cortex, Primary somatosensory (S1) cortex, Primary visual (V1) cortex, Superior temporal cortex, Ventrolateral prefrontal cortex, Primary motor (M1) cortex (Supplemental Fig. S1). Independent temporal comparisons were performed without aggregations of multiple regions. For differential expression between regions, all samples regardless of age were used.

Differential gene expression analysis was performed using voom (Law et al. 2014) as it has been implemented in the R package limma (version 3.46.0). P-values were corrected for multiple testing using the Benjamini-Hochberg’s method (Benjamini and Hochberg 1995). A feature was considered to be differentially expressed when the fold change between the groups compared was higher than 1.5 and the adjusted P-value was below 0.05 or is otherwise stated in figure legends.

### Correlation analysis

Correlation between age and miRNA expression was assessed using spearman correlation and P-values were adjusted using the Bonferroni correction.

### Expression of TE-embedded miRNAs in other tissues

Processed CPM expression data of mature miRNAs in 399 human samples (De Rie et al. 2017 file: human.srna.cpm.txt) was downloaded and log_2_ CPMs of all annotated mature miRNAs overlapping TE annotations were plotted with addition of a pseudo-count of 1.

### miRNA precursor secondary structure analyses

To predict *in silico* miRNA precursor hairpin structures, the DNA sequence of a 200 to 300 bp window around consistent 22bp peaks observed in BAM files was inputted to miRNAfold (Tempel and Tahi 2012; Tav et al. 2016). A stringent threshold of 90% of verified features was used unless otherwise indicated, to ensure only robust hairpins with a very low false positive rate were returned (Tempel and Tahi 2012; Tav et al. 2016).

### AGO-RIPseq data

Processed AGO-RIPseq browser tracks from hESCs-derived neurons was used to confirm TE-embedded annotated and non-annotated miRNAs in the hg38 genome (Petri et al., 2019 files: GSM2850607_Map.CellsAGO2Aligned.out.bw, GSM2850608_Map.CellsAGO2INAligned.out.bw).

### Evolutionary conservation

MULTIZ tracks from the UCSC genome browser were used to determine the presence of non-annotated TE-embedded miRNA sequences in different species (Blanchette et al. 2004; Navarro Gonzalez et al. 2021).

### miRNA target prediction

miRNA target predictions were downloaded from TargetScan Human (Release 8.0) (Agarwal et al. 2015; McGeary et al. 2019; File: Predicted_Targets_Context_Scores.default_predictions.txt). We utilized only the conserved target predictions for conserved miRNAs as defined in TargetScan (Agarwal et al. 2015; McGeary et al. 2019). GO analysis was performed using PantherDB and enrichment was assessed using Fisher’s exact test followed by false discovery rate adjustment using all human genes as background (Mi et al. 2013).

### Brain miRTExplorer application

The Brain miRTExplorer application was implemented in R using the Shiny app package (Chang et al., 2017).

## Availability of data and materials

No new data was generated during the course of this study. Processed data can be interactively visualized using our “Brain miRTExplorer” application at https://tronoapps.epfl.ch/BrainmiRTExplorer/.

## Author contributions

C.P. conceived the study, performed bioinformatic analyses, interpreted the data and wrote the manuscript. S.S. and E.P. performed bioinformatics analyses. C.P. and D.T. edited the manuscript. All authors reviewed the manuscript.

## Acknowledgements

We thank all members of the Trono Lab, Johan Jakobsson and Retha Ritter for helpful and insightful discussions.

