## Supplemental Material for "Transposable elements contribute to the spatiotemporal microRNA landscape in human brain development"

Short title: TE-embedded miRNAs in brain development

### Supplemental Figures

| | | Age (Years) | | | | | | | | | | | | | |
|---|---|---|---|---|---|---|---|---|---|---|---|---|---|---|---|
| | | Childhood (C) | | | | | Adolescence (A) | | | | | | C | A | Total |
| Brainspan Region | Abv. | 1 | 2 | 3 | 4 | 5 | 9 | 12 | 14 | 16 | 19 | 20 | Sum | Sum | Sum |
| Amygdala | AMY | 2 | 1 | 0 | 2 | 0 | 2 | 1 | 0 | 1 | 0 | 1 | 5 | 5 | 10 |
| Cerebellar cortex | CB | 3 | 2 | 0 | 2 | 0 | 2 | 1 | 0 | 1 | 0 | 1 | 7 | 5 | 12 |
| Dorsolateral prefrontal cortex | DFC | 1 | 2 | 0 | 0 | 1 | 1 | 1 | 0 | 1 | 0 | 1 | 4 | 4 | 8 |
| Hippocampus | HIP | 1 | 1 | 1 | 2 | 0 | 2 | 1 | 1 | 0 | 0 | 1 | 5 | 5 | 10 |
| Inferior temporal cortex | ITC | 2 | 2 | 1 | 2 | 1 | 2 | 1 | 1 | 1 | 1 | 1 | 8 | 7 | 15 |
| Medial prefrontal cortex | MFC | 2 | 2 | 1 | 0 | 1 | 1 | 0 | 0 | 1 | 1 | 1 | 6 | 4 | 10 |
| Mediodorsal nucleus of the thalamus | THA | 1 | 1 | 1 | 1 | 1 | 0 | 0 | 1 | 1 | 0 | 1 | 5 | 3 | 8 |
| Orbital prefrontal cortex | OFC | 1 | 2 | 1 | 1 | 0 | 1 | 1 | 0 | 1 | 1 | 1 | 5 | 5 | 10 |
| Posterior inferior parietal cortex | IPC | 1 | 0 | 1 | 2 | 1 | 1 | 0 | 0 | 1 | 1 | 1 | 5 | 4 | 9 |
| Primary auditory (A1) cortex | A1C | 1 | 1 | 0 | 1 | 0 | 1 | 1 | 1 | 1 | 1 | 1 | 3 | 6 | 9 |
| Primary motor (M1) cortex | M1C | 2 | 1 | 1 | 2 | 1 | 1 | 1 | 0 | 1 | 1 | 1 | 7 | 5 | 12 |
| Primary somatosensory (S1) cortex | S1C | 2 | 1 | 1 | 2 | 1 | 1 | 1 | 0 | 1 | 1 | 1 | 7 | 5 | 12 |
| Primary visual (V1) cortex | V1C | 1 | 1 | 1 | 2 | 1 | 1 | 1 | 1 | 1 | 1 | 1 | 6 | 6 | 12 |
| Striatum | STR | 2 | 1 | 1 | 1 | 1 | 1 | 0 | 1 | 1 | 0 | 1 | 6 | 4 | 10 |
| Superior temporal cortex | STC | 2 | 2 | 1 | 2 | 1 | 2 | 1 | 0 | 1 | 1 | 1 | 8 | 6 | 14 |
| Ventrolateral prefrontal cortex | VFC | 2 | 2 | 1 | 1 | 1 | 2 | 1 | 0 | 1 | 1 | 1 | 7 | 6 | 13 |
| | | | | | | | | | | | | Sum | 94 | 80 | 174 |
| | | | | | | | | | | Forebrain Sum | | | 66 | 58 | 124 |

**Figure S1. Summary of samples analysed from the Brainspan Atlas of the Developing Human Brain.** Table highlighting the number of samples per brain region (forebrain regions highlighted in blue) and age. Red indicates no samples for that region and age, orange indicates between one and two samples for that region and age and green indicates three or more samples.

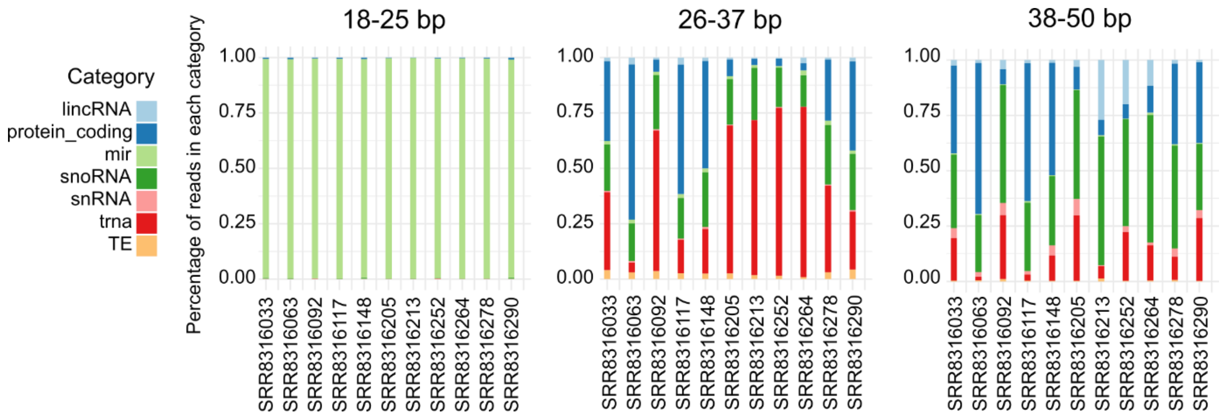

**Figure S2. Different read lengths enrich for different small RNA moieties.** Stacked bar charts indicating the percentage of 18-25bp, 26-37bp or 38-50bp reads overlapping different annotated genomic features for samples from the hippocampus (HIP).

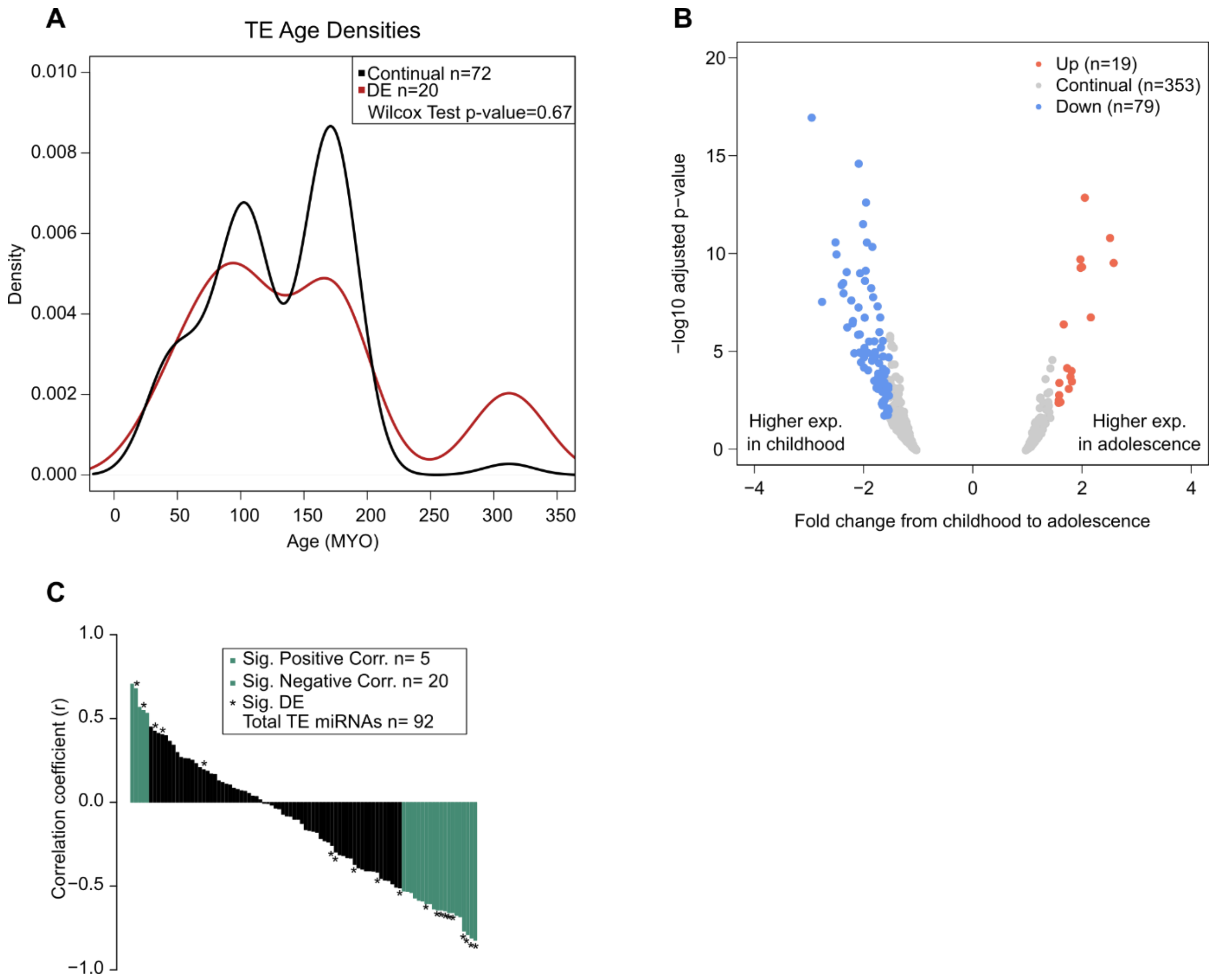

**Figure S3. Differentially expressed TE-embedded miRNAs are a similar age to those continually expressed, their behaviour is similar to non-TE-embedded miRNAs and their expression correlates with donor age.** (*A*) Density plot of TE age in million years old (MYO) of continually and differentially expressed TE-embedded miRNAs. (*B*) Volcano plot highlighting non-TE-embedded miRNAs significantly differentially expressed (adjusted *P-value ≤ 0.05, 1.5-fold change*). (*C*) Bar chart indicating the spearman correlation coefficient of TE-embedded miRNA expression with donor age. Significance was determined as *P-value ≤ 0.05.* * denotes TE-embedded miRNA which are also differentially expressed in Fig. 2A. See also Supplemental Table S2.

### Supplemental Table Descriptions

**Supplemental Table S1 –** Expression of all TE-embedded miRNA and non-TE-embedded miRNA loci in counts per million (CPM) in childhood and adolescence, alongside their differential expression between childhood and adolescence. TE age as defined in DFAM is also shown. The first tab represents the collated analysis on all forebrain samples and the adjusted *P-value* is shown and subsequent tabs provide differential expression analyses showing the non-adjusted *P-value* due to low sample numbers (see Fig. S1 for sample information).

**Supplemental Table S2 –** Spearman correlation analysis of TE-embedded miRNA expression to donor age in collated forebrain samples.

**Supplemental Table S3 –** Information about miRNAs embedded in two individual TEs and their respective orientations.

**Supplemental Table S4 –** Predicted genic TE-embedded miRNA targets from TargetScan. Only conserved miRNAs and conserved targets are included and columns are as defined in Agarwal et al. 2015 and McGeary et al. 2019. File used: Predicted_Targets_Context_Scores.default_predictions.txt.

### Supplemental Acknowledgements

Submission of BrainSpan Atlas of the Developing Human Brain data (phs000406.v2.p1) to dbGaP was provided by Dr. Nenad Sestan. Collection of the data and analysis was supported by grants from the National Institutes of Health (MH089929, MH081896, and MH090047). Additional support was provided by the Kavli Foundation, a James S. McDonnell Foundation Scholar Award, NARSAD, and the Foster-Davis Foundation.
